# Microbiota discovered in scorpion venom

**DOI:** 10.1101/2025.07.03.662918

**Authors:** Barbara Murdoch, Adam J. Kleinschmit, Carlos E. Santibáñez-López, Matthew R. Graham

## Abstract

With low nutrient availability and presence of numerous antimicrobial peptides, animal venoms have been traditionally considered to be harsh sterile environments that lack bacteria. Contrary to this assumption, recent studies of animal venom and venom-producing tissues have revealed the presence of diverse microbial communities, warranting further studies of potential microbiota in other venomous animals. In this study we used 16S rRNA amplicon sequencing to elucidate whether scorpion venom contained bacteria, to characterize the bacterial communities, and determine if venom microbiomes differed across geologically complex geographic locations. Our study compares the venom microbiome of two scorpion species, sampled from sites between the Mojave and Great Basin deserts, *Paruoctonus becki* (family of Vaejovidae) and *Anuroctonus phaiodactylus* (family of Anuroctonidae), and represents the first assessment of microbial diversity ever conducted using the venom secretion itself, rather than the venom-producing organ and its surrounding tissues.

## Introduction

Microbial communities play essential roles in a wide array of host physiological processes, including digestion [1, 2], reproduction [3], and the suppression of other symbionts [4]. Despite their importance, the diversity and potential functions of microbes within animal venom systems remain largely unexplored e.g., [5]. Animal venom, a complex cocktail of inorganic and organic compounds, serves crucial ecological functions in predator deterrence e.g., [6], subduing prey, and even pre-digestion e.g., [7], with some evidence suggesting a role in mating e.g., [8]. Traditionally, venom has been considered sterile. More recently, studies have begun to unveil the presence of microbiomes within snake and spider venom systems, suggesting a potential influence on infections associated with envenomations [9, 10]. This emerging understanding of venom microbiomes underscores the relevance of studying microbial diversity in other venomous animals, such as those found in scorpions.

Scorpions represent an ancient and diverse lineage comprising over 2,800 extant species distributed across nearly all terrestrial habitats and are renowned for their potent venom and remarkable resilience to extreme environments. While the majority of scorpion research has focused on systematics e.g., [11] venom composition and function (reviewed in [12, 13]), and evolutionary origins [14, 15], recent efforts have begun to explore their associated microbiomes [16–18]. Early studies targeting the 16S rRNA gene in the gut microbiota of 24 scorpion species revealed significant bacterial diversity across five scorpion families [16, 17]. More recently, our team’s work has demonstrated a rich diversity of bacteria, including novel phylotypes of class Mollicutes, within the telson of *Hadrurus arizonensis* and *Smeringurus mesaensis* [18]. Although these prior studies assessed microbes of the gut and telson (where venom production occurs), so far none has characterized the venom microbiota, as this study does.

Given that scorpion stings pose a significant global health concern, that secondary infections can complicate envenomation cases [19], and given the vast therapeutic potential of compounds stemming from scorpion venom [20], understanding the microbial communities associated with scorpion venom systems is crucial. While previous studies have identified bacterial genes within the telson and venom glands, the presence and diversity of microorganisms directly within the venom secretion remain unknown. To address this gap, we conducted the first assessment of microbial diversity in scorpion venom using two New World scorpion species from distinct families: *Paruroctonus becki* (Gertsch and Allred, 1965) of family Vaejovidae and *Anuroctonus phaiodactylus* (Wood, 1863) of Anuroctonidae. Both species are distributed across the Mojave and Great Basin deserts but have contrasting ecological preferences: *P. becki* is primarily a psammophilous species often inhabiting alkali-sink environments [21], whereas *A. phaiodactylus* is a pelophilous species burrowing in sedimentary hillsides [22] (Fig. 1). While the telson microbiome of a vaejovid species (*Smeringurus* mesaenesis (Stahnke, 1957)) has been previously investigated [18], this study focuses on the venom microbiome of a different genus within this family (*Paruroctonus*) and provides the first exploration of the microbiome in a member of the Anuroctonidae family. Importantly, as far as we know, our data for both species represents the first assessment of microbial diversity ever conducted using the venom secretion itself, rather than the entire telson.

**Figure 1.**
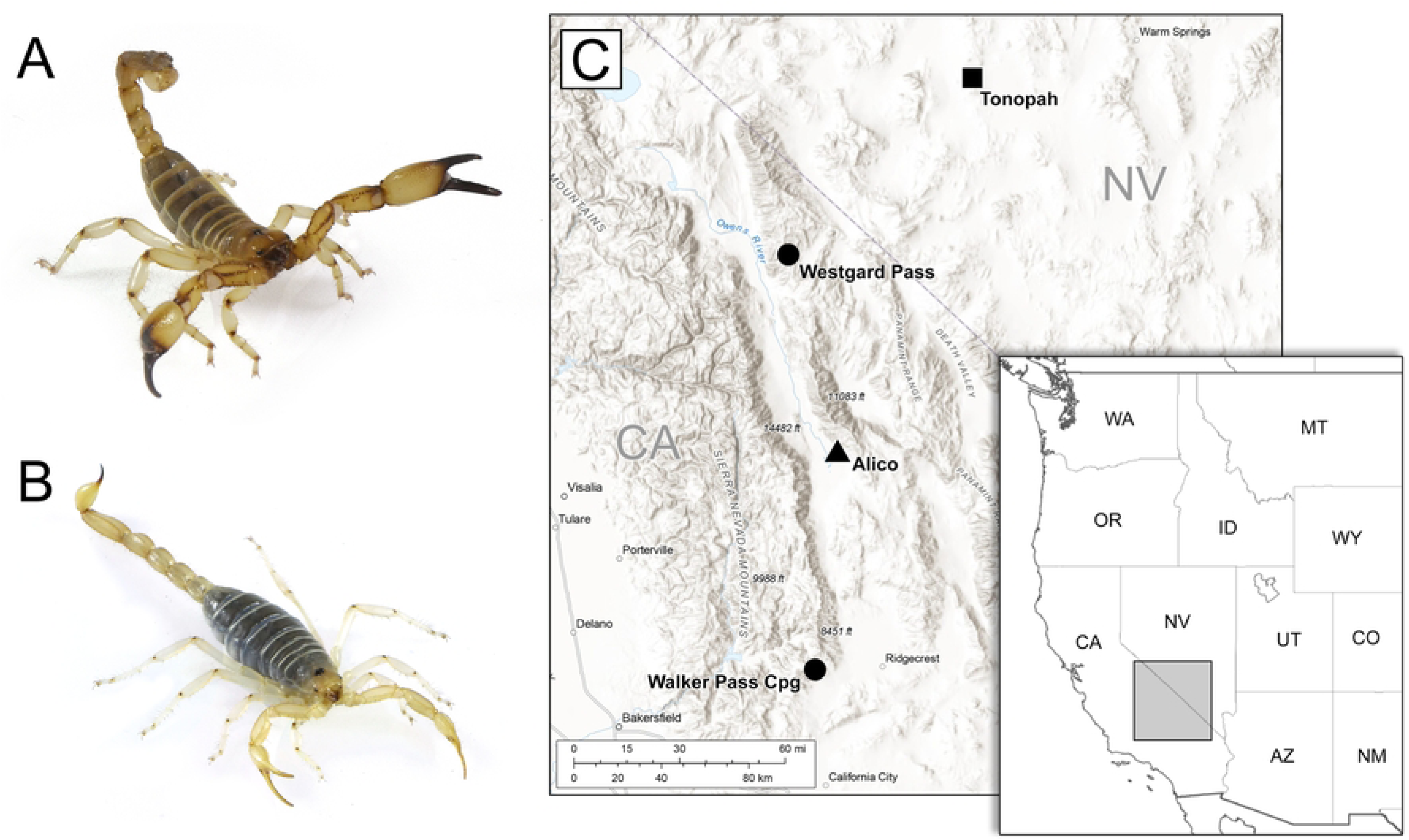
Images of A) *Anuroctonus phaiodactylus* and B) *Paruroctonus becki*. C) Topographic map with regional inset map depicting the distribution of collection sites in California and Nevada. Circles mark individual sample locations where only *A. phaiodactylus* was collected, the triangle is where only *P. becki* was collected, and both species were collected from the site marked with a square.

## Materials and methods

### Scorpion and venom collection

Scorpion collection – We conducted nocturnal field collections using ultraviolet light [23] to obtain 16 *A. phaiodactylus* and 20 *P. becki* specimens from four distinct localities (Permit Number - GW-210220004-21022-001, California Department of Fish and Wildlife). These sites form a latitudinal transect across the transitional ecotone between the Mojave and Great Basin deserts (Fig 1), a region encompassing the Eastern California Shear Zone (ECSZ). The ECSZ, a complex system of active dextral strike-slip faults situated east of the San Andreas Fault, is recognized as a significant geomorphological driver of regional biodiversity patterns. Its long-term influence on topography, hydrology, and habitat fragmentation has been hypothesized to shape evolutionary and ecological processes in various taxa, including *P. becki* [24] and *Aphnopelma* tarantulas [25].

Venom collection – Prior to venom collection, scorpions were sterilized in a laminar flow hood under ultraviolet light for 15 minutes and the telson was additionally sterilized with 70% ethanol, then air dried. Three strategies were used for venom collection. First, scorpions were agitated while enclosed in a sterile plastic bag; venom collection occurred after stinging through the bag onto its outer surface. In the next two strategies, with their aculeus placed into a sterile 1.5 ml Eppendorf tube through ethanol-sterilized parafilm, the scorpion was physically agitated with sterile forceps to elicit venom release. The final strategy involved electrical stimulation using a square wave stimulator of the telson set at 3 events per second, with the duration set to 25 milliseconds and an amplitude of 4 volts. The venom was spun down and DNA was extracted immediately or after storage at - 20°C. Controls included sterile scorpion tails or telsons that were washed in 300 µl of Tissue and Cell Lysis Buffer with 50 µg Proteinase K, collected into sterile tubes.

### DNA extraction from scorpion venom

DNA was extracted using the methods from Shimwell et al., 2023 [18]. Briefly, a modified Master Pure DNA Purification (Biosearch Technologies) protocol was used to extract DNA from the scorpion venom. Venom volumes ranged from about 3 µl to 20 µl per sample. Venom was thawed on ice, mixed with 300 μl Tissue and Cell Lysis Buffer and 50 μg Proteinase K. Samples were resuspended and incubated overnight in a 56°C shaking water bath. Samples were iced and mixed vigorously with 175 μl of MPC Protein Precipitation Reagent, and centrifuged at 10,000 x g for 10 minutes at 4°C. Supernatants were transferred to fresh tubes and mixed with one volume of cold -20°C isopropanol plus 400 ng glycogen and stored at -20°C for 60 minutes to overnight. Samples were warmed to 4°C and centrifuged at 10,000 x g for 10 minutes at 4°C to pellet the DNA. Supernatants were decanted, and the pellets were washed twice with 70% ethanol. The pellets were air dried for 30 minutes and resuspended in 40 μl of TE buffer (10 mM Tris, 1 mM EDTA, pH 8).

### Prescreen for 16S rRNA gene from scorpion venom

Venom was prescreened for the 16S rRNA gene indicative of the presence of bacteria [26] using PCR with the 27FHT (5’AGR GTT TGA TYM TGG CT3’) and 1492RHT (5’ GGY TAC CTT GTT ACG3’) primers. Cycling conditions were as follows: 1 cycle 96°C, 5 minutes; 35 cycles 96°C 1 minute, 58°C 1 minute, 72°C 1 minute 50 seconds; 1 cycle 72°C 10 minutes. Final concentrations were 10 - 50 ng DNA template, 0.5 μM forward and reverse primers, 200 μM dNTPs, 2.5 units Taq DNA polymerase, 20 mM Tris-HCl (pH8.4), 50 mM KCl buffer, 1.5 mM MgCl.

### Illumina sequencing

Illumina sequencing was performed at the Microbial Analysis, Resources, and Services Center, University of Connecticut. The V4 hypervariable region of the 16S rRNA gene was amplified using the 515F (5’-GTGCCAGCMGCCGCGGTAA -3’) and the 806R (5’GGACTACHVGGGTWTCTAAT-3’) primer set and sequenced with Illumina adapters and dual 8 basepair indices [27] on the MiSeq using v2 (2 × 250 bp) paired-end sequencing platform (Illumina).

## Bioinformatics analysis

### *P. becki* and *A. phaiodactylus* venom microbiome analysis

Demultiplexed fastq sequences were obtained after sequencing from the sequencing center. The quality of the raw sequence reads were assessed by a combination of FastQC [28] and MultiQC [29] software. Bioinformatics analysis was performed using the QIIME2 pipeline version 2023.5 [30]. Descriptive statistics and sequence quality information were further reviewed after demultiplexed raw FASTQ files were imported into QIIME2. Based on the raw sequence quality statistics, raw reads were truncated at bp position 191 to remove low quality base calls and the DADA2 software package was used to trim (191 bp), filter, merge paired reads, remove chimeras, and dereplicate sequences [31]. To reduce noise from singletons and rare amplicon sequence variants (ASVs), ASVs that composed of <0.1% of smallest sample (∼9,700 reads) were filtered. Alpha rarefaction plotting was used to assess if there was sufficient sequencing depth to represent sample diversity. A sampling depth of 2,100 sequences was used for alpha and beta diversity analysis. The resulting rarefied feature table and ASVs were used to generate a phylogenetic tree. ASV taxonomy was assigned using a pre-trained Naive Bayes taxonomic classifier specific to the V4 region (515F-806R) of the 16S rRNA gene using the Greengenes 2 database [32].

Amplicon data were transformed using R package phyloseq [33] to generate taxon abundance visuals and beta diversity visuals in R (version 3.3.1). Beta diversity was computed using the Weighted Unifrac distance and visualized through Principal Coordinate Analysis (PCoA).

Core Microbiome - A composite 2,100-rarefaction depth *P. becki* and *A. phaiodactylus* feature table was split by scorpion species using QIIME2. The composite and split dataset core microbiomes were analyzed independently using phyloseq (v1.34.0) [33] and microbiome (v1.23.1) [34] software packages for R programing language (v4.3.0). These software packages can be accessed via https://github.com/joey711/phyloseq and https://github.com/microbiome/microbiome-respectively. Differential abundance was analyzed using ANCOM-BC [35].

#### *P. becki* exterior control vs. venom microbiome analysis

A large subset of the *P. becki* exterior control samples exhibited low quality Phred scores at the 3’ ends, thus due to a lack of overlap between forward and reverse reads after truncation at 80bp, only forward sequences were used for exterior control vs. venom analysis. After forward single-end sequence truncation at 80bp of samples used in this analysis via DADA2 in the QIIME2 pipeline outlined above, the samples were rarefied to a depth of 17,943 sequences.

### Statistical analysis

Differences between alpha diversity indices were tested using the Kruskal–Wallis test (QIIME2). Beta diversity metric distance was statistically tested by non-parametric multivariate ANOVA (PERMANOVA) with 999 permutations using QIIME 2 software package.

## Results

We used Illumina 16S rRNA amplicon sequencing to determine whether scorpion venom contains bacteria, and to characterize the bacterial communities. We sampled venom from two scorpion species, *Anuroctonus phaiodactylus* (n=16 scorpions) and *Paruroctonus becki* (n=20 scorpions) collected from various regions encompassing the Eastern California Shear Zone in California and Nevada, that is known as a driver of regional biodiversity (Fig 1). Some samples were run in duplicate on separate sequencing runs, *A. phaiodactylus* (n=31 samples sequenced) and *P. becki* (n=23 samples sequenced). After processing for quality filtering, trimming, chimera removal, *etc*., we generated 2,528,745 16S rRNA paired-end sequence reads, 1,679,193 from *A. phadioactylus* and 849,552 from *P. becki*.

To validate that our sequence reads were in fact sourced from the venom rather than the environment, we compared the bacterial sequences detected in the venom of *P. becki* versus paired exterior controls (n= 6 for each). Beta diversity metrics visualized via Bray Curtis principal component analysis plots showed differences in the microbial composition of the exterior controls compared to the venom (p<0.006; S1 Fig). Similar results for beta diversity were reached using weighted Unifrac (p<0.036) and Jaccard (p<0.004), but not unweighted Unifrac (p<0.118; S1 Fig). Collectively, these results suggest that the amplicon sequence variants (ASVs) identified in the venom were not due to environmental contamination and that the differences detected were driven largely by differential relative abundance.

Alpha rarefaction plotting was used to assess a sufficient sequencing depth to represent sample diversity (S2 Fig). To include as many samples as possible while covering the majority of the detected diversity, we chose a depth of 2,100 sequence reads for downstream assessment of ASVs, including alpha and beta diversity. After processing, these sequence reads were assigned to 1,841 total ASVs; 159 ASVs (8.6%) were shared between the venom of both scorpion species. Of the 1,325 total ASVs for *A. phaiodactylus,* 88% were unique, whereas 69% (total 516 ASVs) were unique to *P. becki* (S3 Fig).

### Bacterial diversity of scorpion venom

The ASVs were assigned taxa using the Greengenes 2 database (McDonald et al., 2023). The most relatively abundant phyla for *A. phaiodactylus* vs *P. becki* respectively, were Pseudomonadota (58% vs 77%), Actinobacteriota (15% vs 3%), Bacillota (14% vs 6%), Bacteroidota (10% vs 12%), and “Others” that includes phyla each with reads of less than 1% (3% vs 2%; Fig 2A, B). A heatmap at the family level shows similar relative abundances between samples for *A. phaiodactylus* and *P. becki* except for the 4 and 7 taxa at the top of the heatmap of columns A and D for *phaiodactylus* and *P. becki,* respectively, which appear to have lower relative abundances (Fig 2C). These samples were collected from the regions of Tonopah, NV, for *A. phaiodactylus* and Alico, CA, for *P. becki,* suggesting a possible link between a region and the venom microbiome.

**Figure 2.**
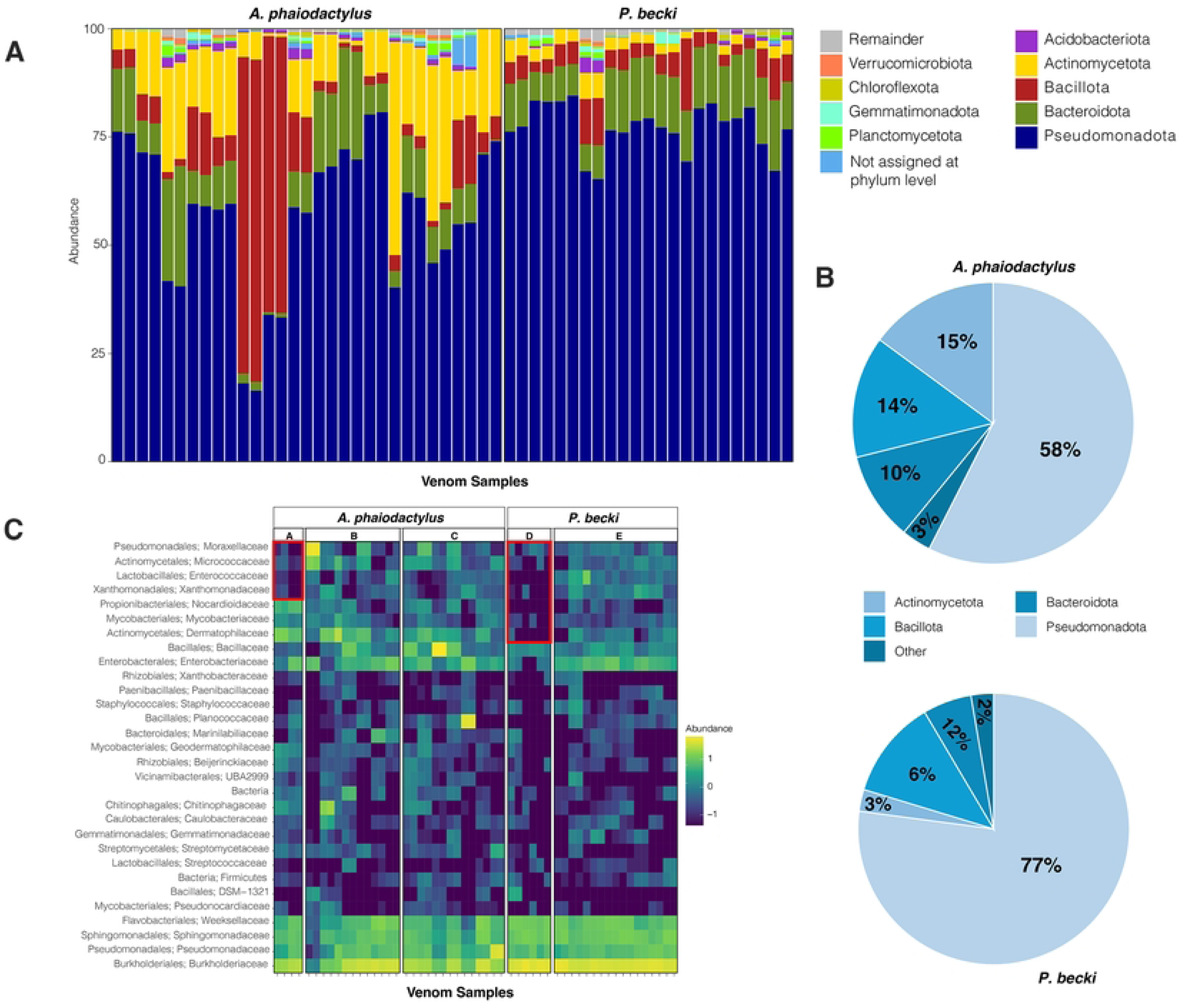
Taxonomic relative abundance profiles at the phylum and family level for *A. phaiodactylus* vs. *P. becki* venom microbiome samples. A) 100% stacked bar plot of top 10 phyla; B) Pie charts summarizing percentage abundance for *A. phaiodactylus* and *P. becki*. C) Heatmap of top 10 family taxa. Geographical regions include: A, E – Tonopah, NV; B – Walker Pass Campground, CA; C – Westgard Pass, CA; D – Alico, CA.

For the genera in each venom microbiome (Fig 3A), we found a higher percentage of reads in *P. becki* compared to *A. phaiodactylus* for *Ramlibacter* (47% vs 24%), *Sphingomonas* (12% vs 7%), and *Chryseobacterium* (11% vs 7%), and similar percentages for *Pseudomonas* (8% vs 9%) and *Proteus* (2% vs 3%; Fig 3B). Additional genera seen in *A. phaiodactylus* above 1% of the total reads included *Bacillus* (6%), *Acinetobacter* (5%), *Psychrobacillus* (4%), *Ectobacillus* (2%), and *Nocardioides* (1%; Fig 3B). Although *Psychrobacillus* was not detected in *P. becki,* these other genera were, but at relative abundances lower than 1% (Fig 3B). For *P. becki* and *A. phaiodactylus* respectively, 20% and 32% of all reads represented relative abundances of less than 1% (Fig 3B). To assess differentially enriched and depleted microbiota, we performed an analysis of compositions of microbiomes with bias correction (ANCOM-BC, Fig 3C). In *P. becki,* our analyses showed enrichment for *Ramlibacter, Chryseobacterium*, and *Sphingomonas* (p<0.05), in addition to *Leuconostoc*, *Flavobacterium*, *Atopostipes*, and the family of *Gaiellaceae* (p<0.01). In contrast *Nocardioides* and members of the family *Dermatophilaceae* were depleted in *P. becki* relative to *A. phaiodactylus* (p<0.05; Fig 3C).

**Figure 3.**
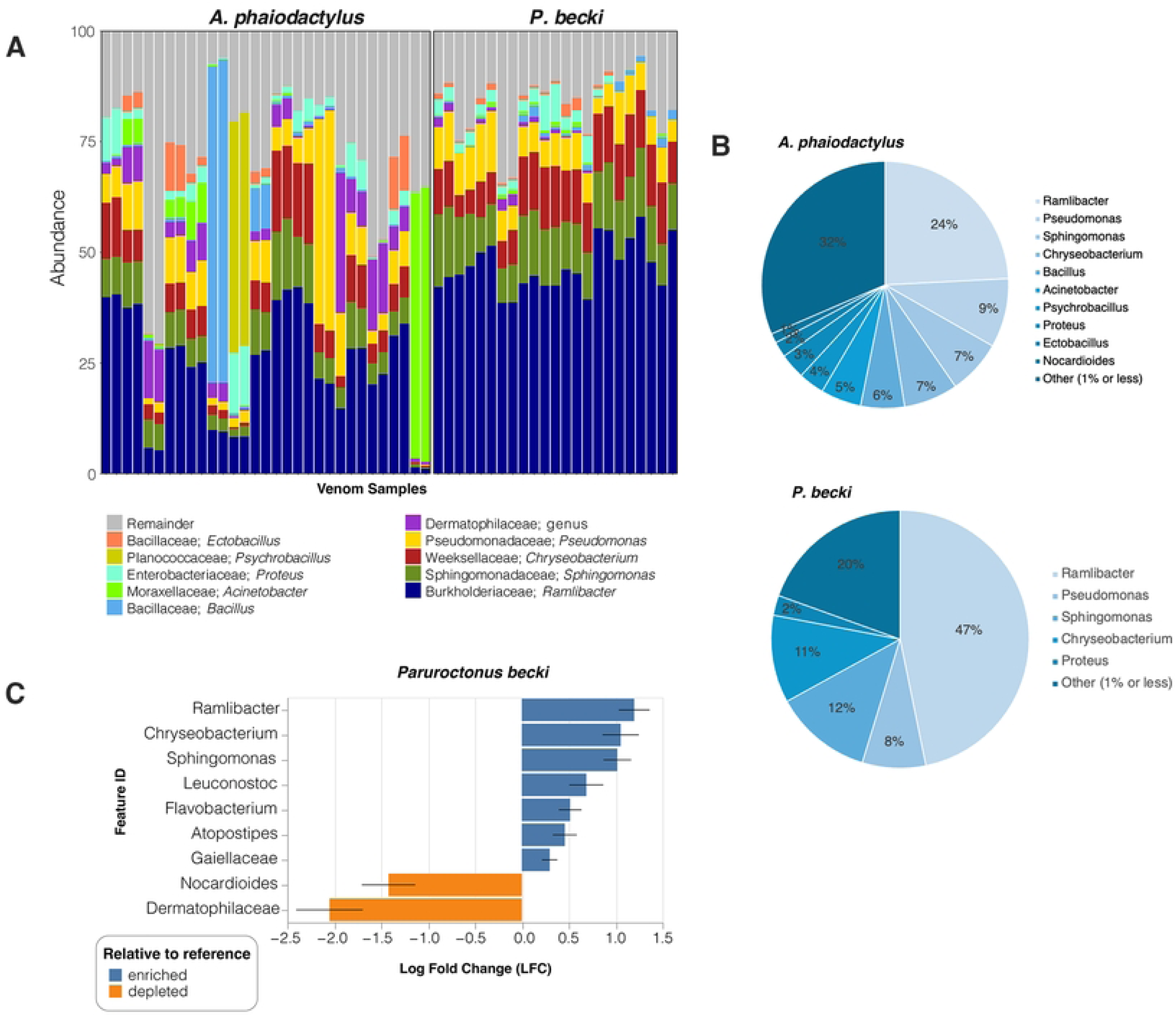
Taxonomic profiles at the genera level for *A. phaiodactylus* vs. *P. becki* venom microbiome samples. A) 100% stacked bar plot of top 10 genera; B) Pie charts summarizing percentage abundance for all *A. phaiodactylus* and *P. becki* samples. C) ANCOM-BC differential abundance log fold change (LFC) for genera of *A. phaiodactylus* and *P. becki*. Reported taxa exhibit significantly different abundances (Holm adjusted p<0.05) between the two scorpion species.

### The core venom microbiome of *A. phaiodactylus* and *P. becki*

We further investigated the taxonomic composition of the venom microbiome in the two scorpion species by focusing on identifying their core microbial constituents. The core microbiome was defined using two conservative criteria: (1) a relative abundance threshold of 0.01% and [30] an occurrence in at least 50% of the samples. These thresholds were selected to strike a balance between capturing ecologically relevant, low-abundance taxa and minimizing noise [36], especially given the rarefaction depth of 2,100 reads. S1 Table shows the number of core microbiome ASVs and their fraction of the total number of ASVs identified at different levels of occurrence. At the ASV level and 50% occurrence threshold, the core microbiome represented a total diversity comprising 1.4% and 4% of the detected genera in *A. phaiodactylus* and *P. becki*, respectively (S1 Table). While the broader microbial communities in the venom were heterogeneous, a relatively stable and tightly aligned core set of microbial taxa emerged independently in both species. ASVs with taxonomy assigned to the genera *Ramlibacter*, *Chryseobacterium*, *Pseudomonas*, and *Sphingomonas* dominated the core microbiome (Fig 4A, B). Of the 159 shared ASVs independent of occurrence between *P. becki* and *A. phaiodactylus* (S3 Fig), 20 formed the shared ASV level core microbiome (≥ 50% occurrence).

**Figure 4.**
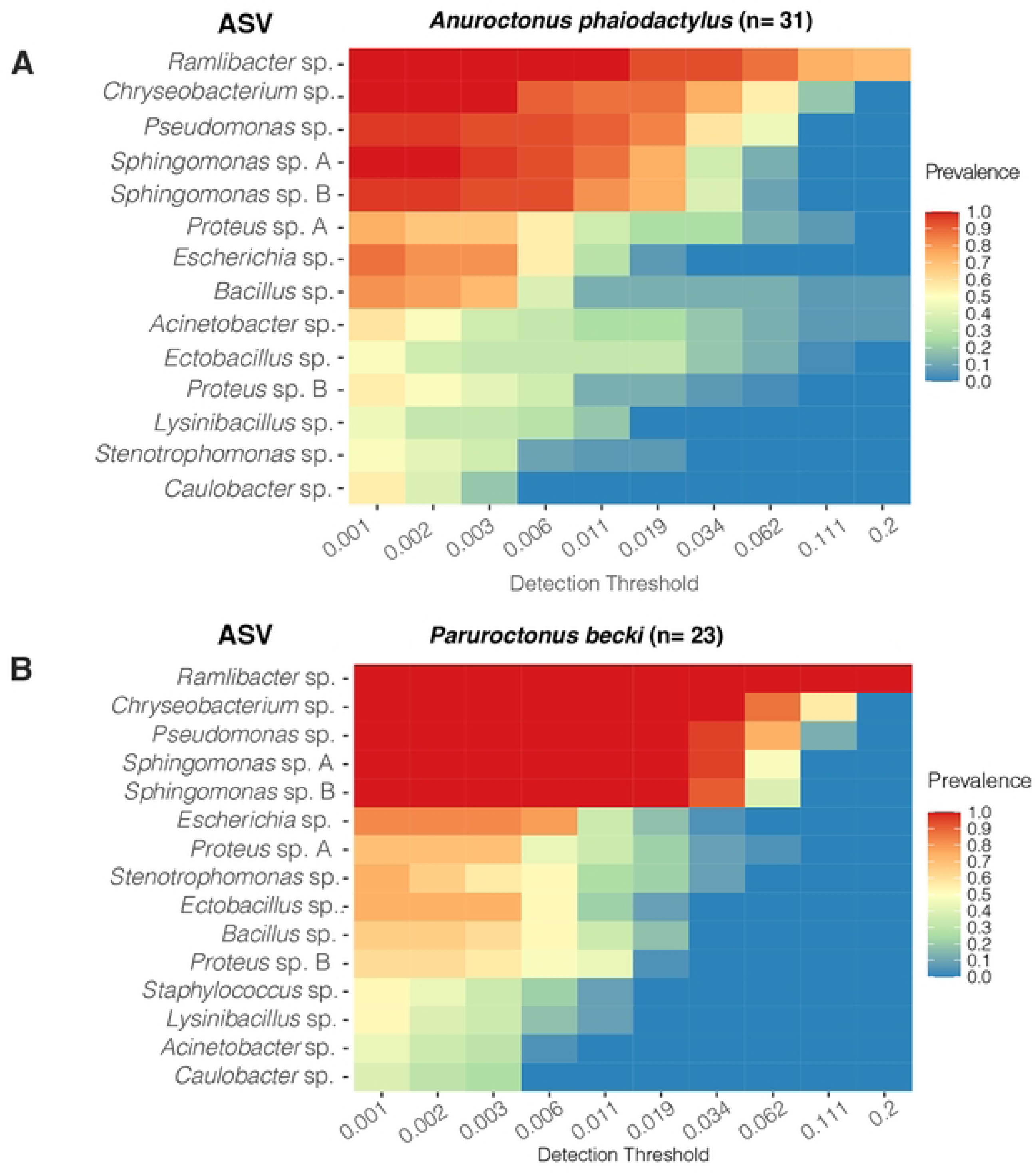
Heatmaps of the abundance–occurrence relationship of the *P. becki* and *A. phaiodactylus* core venom microbial ASVs. A) *A. phaiodactylus*, n=31. B) *P. becki*, n=23. The feature table was rarefied to a depth of 2,100 sequences. Minimal thresholds of 0.01% for relative abundance and 50% for occurrence were applied. Taxonomy naming convention indicates the ASV-associated genus.

### Microbial community diversity in *A. phaiodactylus* and *P. becki*

Alpha diversity was performed to determine the within species taxa variation in each scorpion venom bacterial community. For the venom taxa found in *A. phaiodactylus* and *P. becki,* respectively, richness was assessed using Observed taxa (mean ± standard error) 102 ± 11; 56 ± 5, and ACE diversity 125 ± 16; 60 ± 6, that indicated significant differences in alpha diversity between species for both measures (Fig. 5A, B; S2 Table, p<0.05). Differences in Shannon 4 ± 0.2; 3 ± 0.003 and Simpson (0.8 ± 0.02; 0.8 ± 0.01 diversity, for *A. phaiodactylus* and *P. becki,* respectively, were also statistically significant for the venom microbiome in each scorpion species (Fig 5C, D; S2 Table, p<0.05).

**Figure 5.**
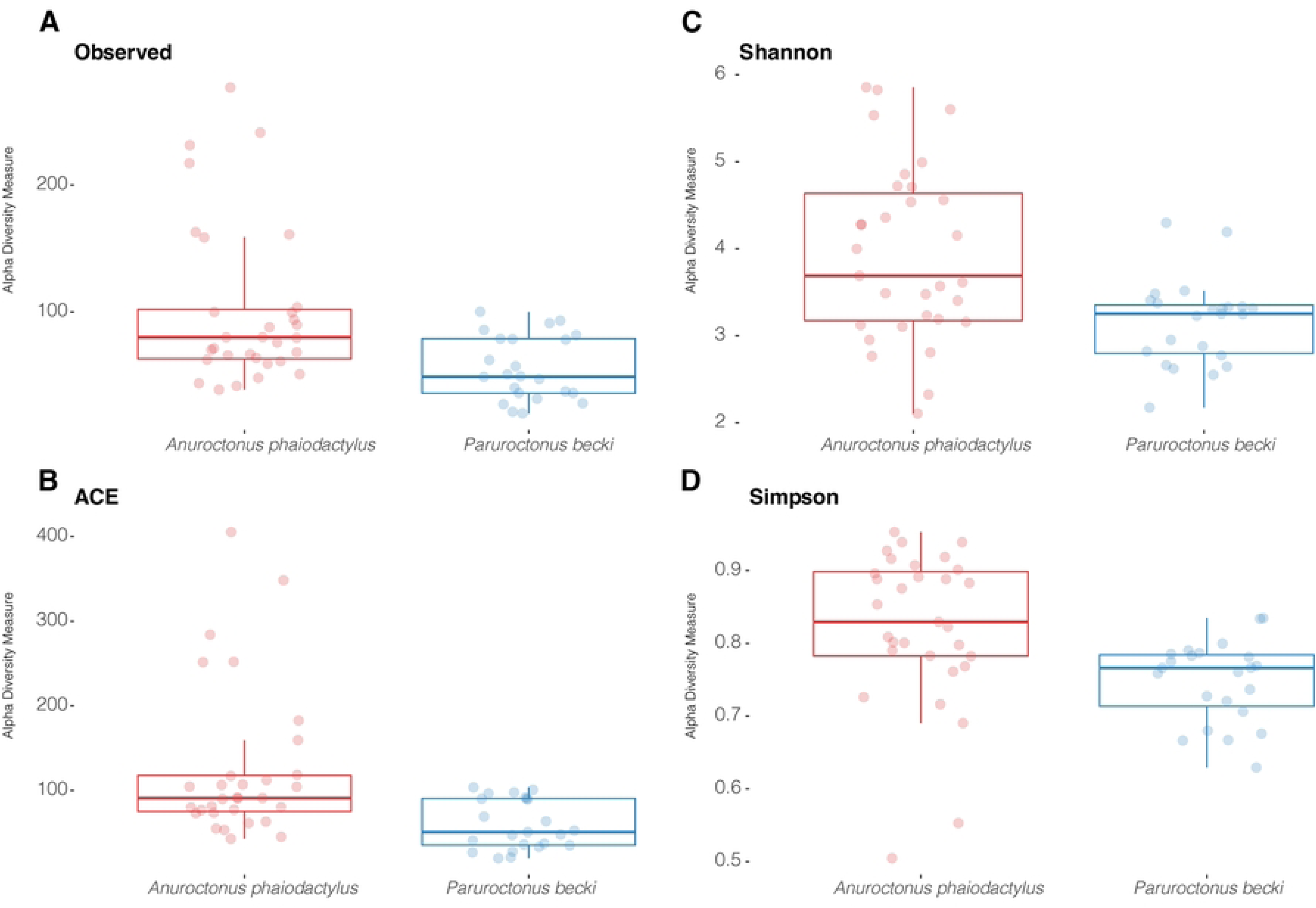
Alpha diversity box plots of *A. phaiodactylus* vs. *P. becki* venom microbiome samples. Species richness was inferred using A) Observed taxa and B) ACE diversity metrics, while C) Shannon, and D) Simpson diversity metrics were used to look at how differentially weighting richness and evenness influenced diversity. Pairwise Kruskal-Wallis statistical testing indicated significance for all measures (p<0.05).

Additional metrics of alpha diversity showed significant differences in the venom microbiota for both scorpion species (S2 Table; p<0.05). For example, for *A. phaiodactylus* and *P. becki,* respectively, Faith’s Phylogenetic Diversity (mean ± standard error) was 9.15 ± 0.78 and 6.48 ± 0.40); Pielou’s Evenness was 0.61 ± 0.02 and 0.5 6 ±0.01.

Beta diversity measures of Bray Curtis, weighted Unifrac, Jaccard, and unweighted Unifrac, all showed statistical significance (Fig 6A-D; PERMANOVA, p<0.001), indicating different microbial communities found within the venom of *A. phaiodactylus* compared to *P. becki*. Measures relying on presence/absence, like Jaccard and unweighted Unifrac, appeared to have some overlap, indicating some similarities, rather than clear distinctions (Fig 6C, D). In contrast, measures relying on relative abundance such as Bray Curtis and weighted Unifrac, appeared more distinct in space (Fig 6A, B). These indices suggest that the observed beta diversity metrics are driven by relative abundance rather than presence/absence.

**Figure 6.**
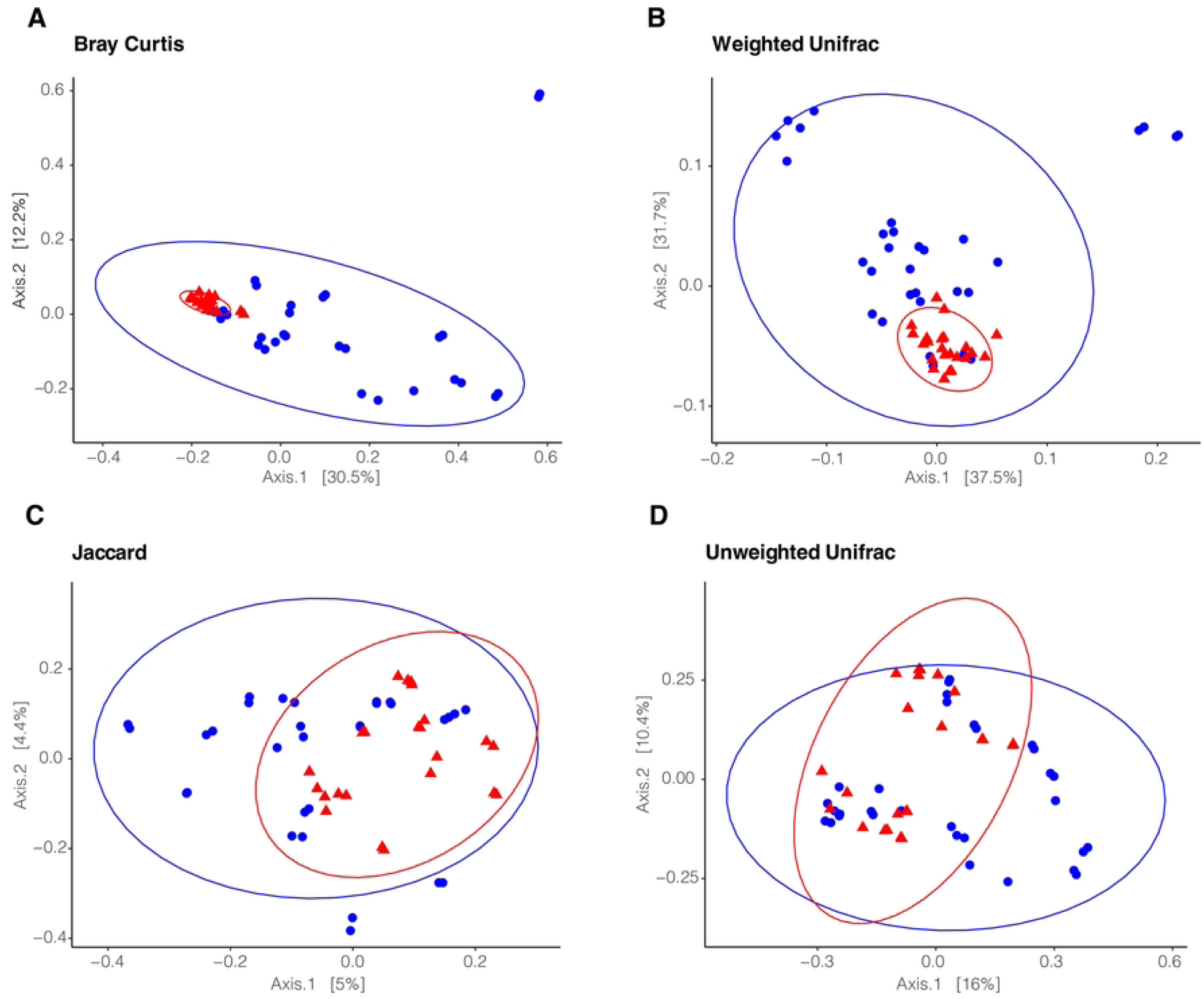
Beta diversity 2D PCoA ordination plots of *A. phaiodactylus* vs. *P. becki* venom microbiome samples. Beta diversity was examined using A) Bray Curtis, B) Weighted Unifrac, C) Jaccard, and D) Unweighted Unifrac. Pairwise statistical testing for all distance metrics was significant (PERMANOVA, 999 permutations, p<0.001).

### Geographical location affects the venom microbiome

We collected scorpion samples from different locations in California and Nevada (Fig 1) and tested whether geographical location influenced their venom microbiomes.

We found the geographical location to be a significant factor influencing the microbial diversity of both scorpion species (Fig 7). For measures of richness (Observed and ACE) the median in *A. phaiodactylus* was highest in the region of Tonopah, NV, compared to either Walker Pass Campground, CA or Westgard Pass, CA, and Tonopah, NV showed a greater variability in sample diversity (Fig 7A, B). Similar results for richness were seen for *P. becki*, where Tonopah, NV compared to Alico, CA had the highest median and greater variability in sample-to-sample diversity (Fig 7 A, B). For both scorpion species, alpha diversity metrics that account for richness and evenness, like Shannon and Simpson, supported the differential diversity of the Tonopah, NV region compared to the California-based regions (Fig 7 C, D). Except for Observed taxa, all other metrics (ACE, Shannon and Simpson) were significant (Holm adjusted p<0.05). Our results suggest that microbiome diversity can be shaped by geographical location, and are potentially influenced by regional environmental factors, habitat, behavior, or host genetics.

**Figure 7.**
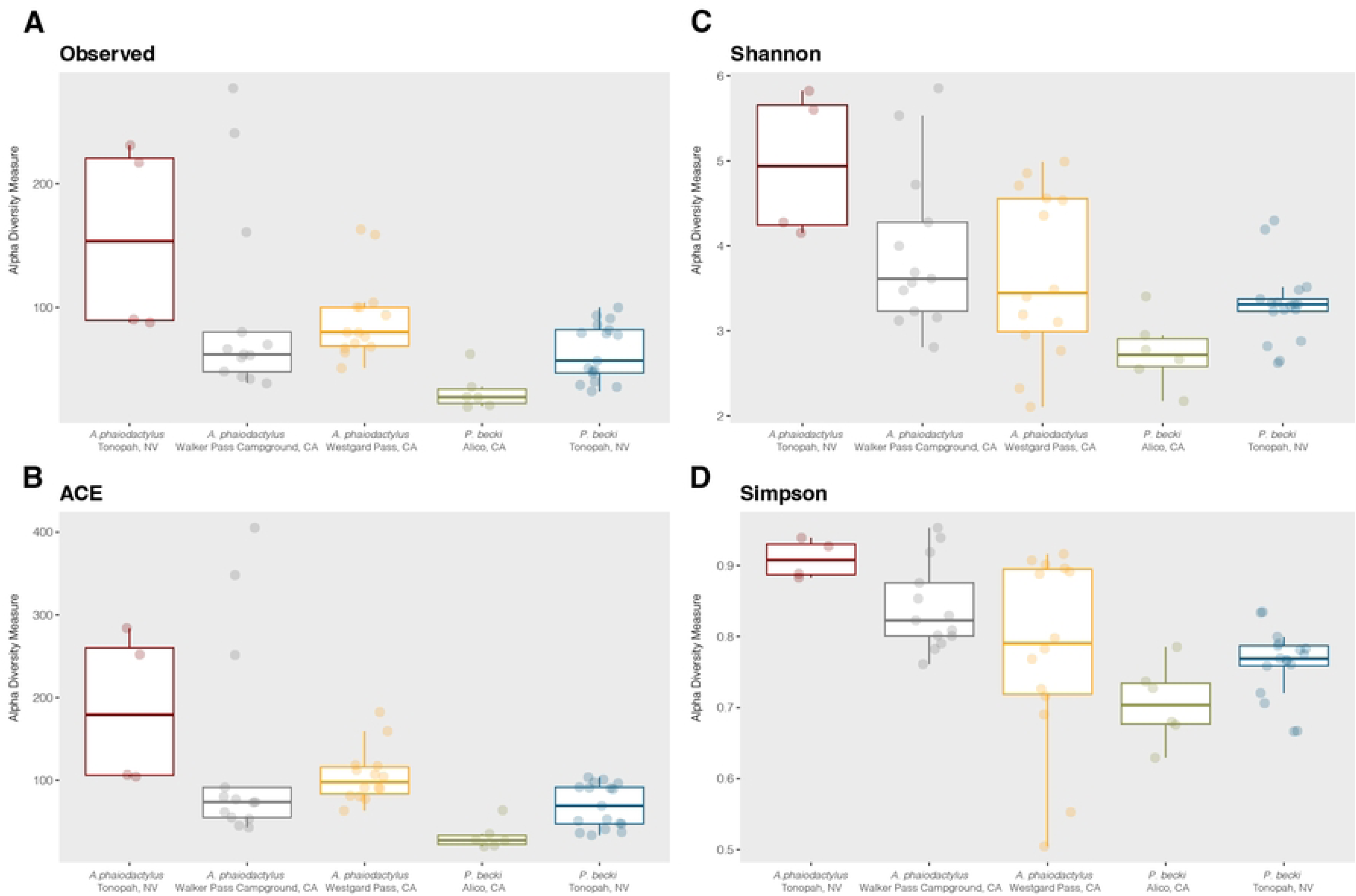
Alpha diversity box plots of *A. phaiodactylus* vs. *P. becki* venom microbiome split by geographical location. Species richness was inferred using (A) Observed taxa and (B) ACE diversity metrics, while (C) Shannon, and (D) Simpson diversity metrics were used to look at how differentially weighting richness and evenness influenced diversity. For *A. phaiodactylus* vs. *P. becki at Tonopah, NV:* Observed, q= 0.24; ACE, q=0.0082; Shannon, q=0.027; Simpson, q=0.0078.

### Differential abundance analysis of functional metabolic pathway potential

We performed a functional profile analysis using PICRUSt2 [37] to predict potential metabolic pathways that operate within the scorpion venom microbiome of *A. phaiodactylus* and *P. becki*. In comparing the abundance of the top 21 KEGG features using *A. phaiodactylus* as a baseline, we found a relatively higher abundance in *P. becki* for bacterial chemotaxis, ABC transporters, two-component system, for the metabolism of glutathione, and metabolism via cytochrome P450 of drugs and xenobiotics (Fig 8A, B). Conversely, several features were more relatively abundant in *A. phaiodactylus* with the most abundant involving fatty acid biosynthesis, terpenoid biosynthesis and benzoate degradation (Fig 8A, B). The overall abundance data for all measured pathways across samples shows two distinct clusters, one for each scorpion species, that are best separated along the PC1 axis which accounts for almost 50% of the variation. (Fig 8C). The density plots along the axes further illustrate this separation indicating distinct distribution peaks for each species (Fig 8C). Heatmap scores for the predicted metabolic pathways from PICRUSt2 analysis that function in the venom microbiome of each scorpion species are shown in S4 Fig. Based on these KEGG features, the data indicate significantly different predicted metabolic/functional profiles for *A. phaiodactylus* compared to *P. becki*.

**Figure 8.**
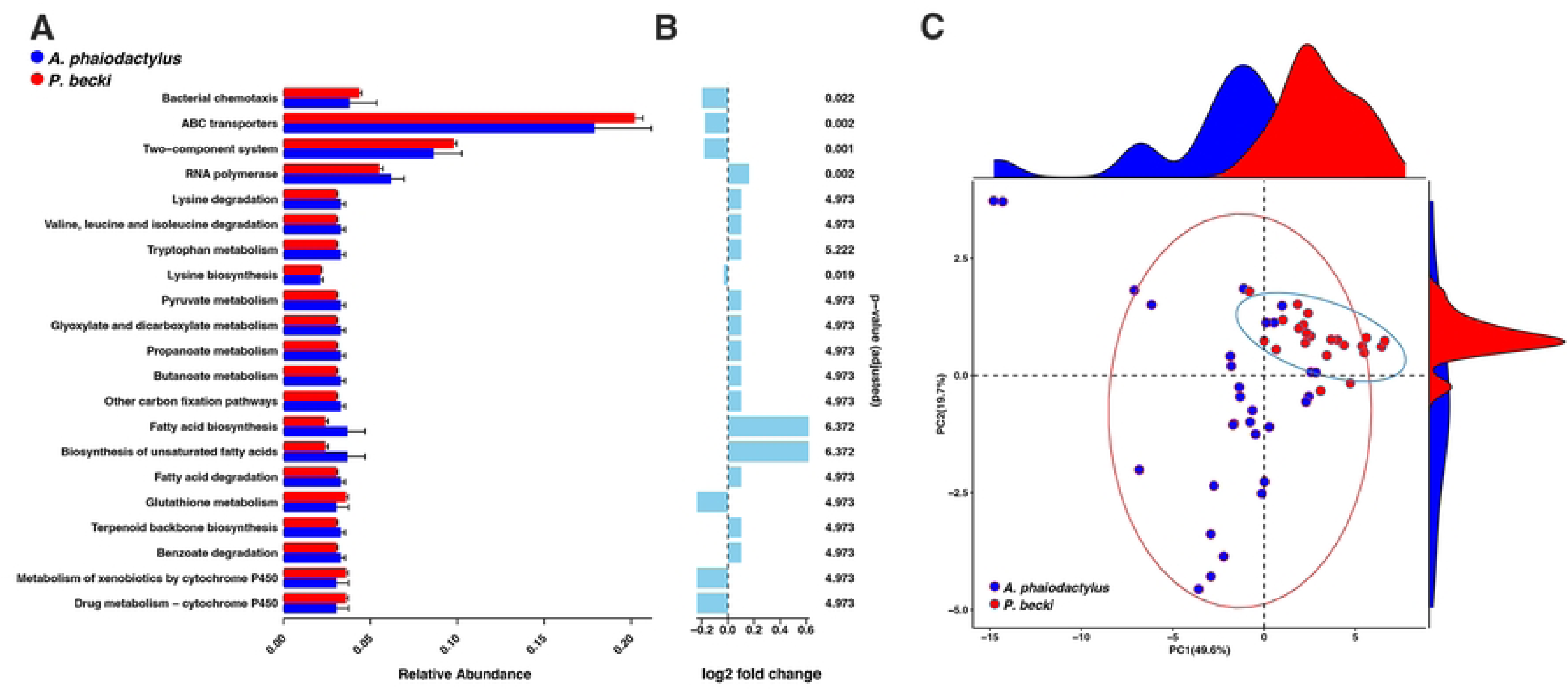
PICRUSt2 analysis of bacterial communities derived from the most abundant reads in the venom of *Anuroctonus phaiodactylus* and *Paruroctonus becki*. A,B) The top 21 KEGG features that were differentially abundant between the two species (ALDEx2_Welch’s t-test, p< 0.01). C) Principal component analysis clustering the functional differences between the bacterial communities in the two scorpion species.

## Discussion

This study, the first of its kind to study scorpion venom microbiota, refutes the notion that venom is a sterile environment devoid of bacteria and corroborates the discovery of microorganisms in the venom of other organisms, like snakes and spiders [9, 10]. Next-generation amplicon sequencing of scorpion venom from *A. phaiodactylus* and *P. becki* revealed a shared core microbiome, but a significant range of diversity within and between species, the latter being driven by differential relative abundance. Further, venom microbiota showed significantly varying alpha diversity by geographical location, with the Tonopah, NV population in the Great Basin desert, being more diverse than the California locations to the south.

Extracting venom for scientific research presents well-documented challenges, as experienced by our team and others [9, 10]. These difficulties stem not only from the need for specialized expertise in safely collecting and handling venomous animals but also from several limiting factors during the extraction process itself. These limitations include the proportion of animals that yield venom, the small volume of venom released per individual—particularly relevant for subsequent DNA extraction—and potential data loss during sequencing and analysis. In our study, we successfully extracted individual venom samples from an average of 92% of our scorpions: 89% (16/18) for *A. phaiodactylus* and 95% (20/21) for *P. becki*. Furthermore, we successfully extracted DNA from all collected venom samples, indicating minimal losses during these initial project phases. Most data losses occurred during the analysis phase. While all *A. phaiodactylus* samples (16/16) advanced through the analysis pipeline, only 70% (14/20) of the *P. becki* samples could be included in the final analysis due to insufficient sequencing reads.

The microbial communities inhabiting metazoans appear to be shaped by a combination of ecological, environmental, and host-specific factors [38]. Environmental context, such as geographic location and habitat type, may influence the presence and composition of distinct microbial taxa, with certain bacteria exhibiting a stronger capacity to adapt to anatomical microhabitats [39]. The study of core taxa may assist in identifying microbial persistence requirements within the habitat [38, 40].

The microhabitat of the venom gland, the site of venom production, may select for bacteria with resilience in adaptive ability and functional features that allow establishment after gaining access from the external environment [9, 41]. These microcosms in venom-associated tissue are likely to be defined by extreme and selective pressures, including the presence of antimicrobial peptides and low nutrient availability, which likely act as environmental filters that favor microbial taxa with resilience and adaptive functional traits. Found to be differentially enriched in the venom microbiota of *A. phaiodactylus* compared to *P. becki,* was the genus *Nocardioides* (family Dermatophilaceae), which can endure extreme environments, and are known for their biotransformation capacity, including pollutant degradation and toxin removal, in addition to production of antimicrobial compounds [42]. Interestingly, antimicrobial compounds in animal venoms are presumed to be made by the host, but the discovery of venom-based microbial communities in this study of scorpions and other studies of snakes and spiders, validates the notion of antimicrobial production by the inherent bacteria [9, 10, 18].

The open structure of the scorpion venom apparatus, particularly the ducts exposed near the tip of the aculeus, may offer an access point for opportunistic environmental microbes. However, the successful colonization of this niche likely requires more than passive entry, as microbes must also withstand the antimicrobial components of the venom and potentially exploit protective or symbiotic mechanisms to persist. One genus that exemplifies such adaptability is *Ramlibacter*, which was found to have a 100% prevalence and accounted for 24% and 47% of genera from *A. phaiodactylus* and *P. becki* scorpion species, respectively. Taxonomically placed within the Pseudomonadota phylum (order Burkholderiales, family Comamonadaceae), *Ramlibacter* is known for its high environmental resilience, including tolerance to desiccation, dormancy under nutrient-poor conditions, and persistence in arid terrestrial environments [43]. These conditions closely align with the arid habitats where the scorpions in this study were collected. Interestingly, *Ramlibacter* has also been reported in the venom gland microbiome of the spider *Steatoda grossa*, a metropolitan species sourced from captive conditions [44], suggesting that this genus may opportunistically colonize venom glands across diverse arthropod hosts. Furthermore, *Ramlibacter* has been detected in a wide range of terrestrial and aquatic environments worldwide increasing the likelihood of colonization of diverse venomous species [44].

In addition to *Ramlibacter*, we observed the complete prevalence and relatively high abundance of *Sphingomonas*, *Chryseobacterium*, *Pseudomonas, and Proteus* across scorpion venom samples. These genera have also been documented in the venom or venom-associated tissues of multiple spider species [10, 41, 45], and *Pseudomonas* showed the highest relative abundance in blood samples from patients following snake bites [46], highlighting a potential pattern of convergence in venom gland-associated microbiota among distantly related venomous metazoans. These findings support the hypothesis that venom glands may serve as selective environments favoring certain taxa with functional traits conducive to survival in such chemically hostile niches.

The potential for microbial ingress through external structures is further supported by examples in other venomous animals. In spiders, opportunistic microbes are thought to colonize the envenomation apparatus via external chelicerae, while in snakes, biofilm formation has been implicated in the oral cavity and venom delivery structures [9]. Similarly, scorpions undergo cleaning behaviors before and after feeding, for example to clean their stingers with their chelicerae, or after a sting by rubbing the stinger on a substrate like sand [47]. These behaviors suggest potential microbial entry into the scorpion venom organ through the oral cavity, the external environment, or through captured prey, either after being stung or after being consumed by the scorpion.

Beyond the most dominant core taxa, our study also identified members of the Enterococcaceae family, that includes the genus *Enterococcus*, at or below 0.2% of total reads for both scorpion species, which has previously been isolated from the venom of the black-necked cobra. Notably, *E. faecalis* strains from that study possessed genes associated with enhanced membrane integrity and antimicrobial resistance, which suggests microbes with characteristics beneficial for the venom environment may be selected for [9]. Collectively, these findings raise thought-provoking inquiries about the potential functional roles and genomic adaptations of scorpion-associated microbial taxa.

Other genera detected in our sample core microbiome, including *Stenotrophomonas*, *Acinetobacter*, *Bacillus*, and *Staphylococcus*, have been previously reported in venom-associated tissues of spiders and marine gastropods [9, 10, 41, 48]. For example, *Stenotrophomonas* spp. have been proposed to interact directly with venom peptides, potentially modifying their biological activity. The detection of Caulobacteraceae taxa in our sample core adds further depth to the taxonomic range of scorpion venom gland-associated microbes also detected in other venom-associated tissues [48], warranting future investigation into their ecological and functional roles. The clinical potential of venom compounds suggests the possibility, that microbes within their unique venom microhabitats may provide the necessary chemical precursors for the production of these therapeutic compounds.

Our unique multi-site sampling of venom microbiomes from *A. phaiodactylus* and *P. becki* offers insights into the influence of geographic and geological factors on microbial communities in venoms. All samples were collected within the tectonically complex Eastern California Shear Zone (ECSZ), an area where significant geological activity [49] is presumed to have driven diversification in both *P. becki* [24] and *Aphonopelma* tarantulas [25]. For *P. becki*, our samples encompass two distinct clades as identified by Graham et al. (2013): the White-Inyo Clade (Alico, CA) and the Great Basin Clade (Tonopah, NV). Similarly, our *A. phaiodactylus* samples were collected from three localities within the ECSZ region: the southern Sierra Nevada (Walker Pass Campground, CA), the White-Inyo Mountains (Westgard Pass, CA), and the Great Basin Desert (Tonopah, NV). Unpublished phylogenomic data (Graham) suggest that these *A. phaiodactylus* sites also represent distinct clades. Interestingly, the Tonopah samples from the Great Basin consistently exhibited the highest venom microbiome diversity for both scorpion species (Fig. 7). This observation aligns with phylogeographical hypotheses suggesting that both *P. becki* and *A. phaiodactylus* underwent recent range expansions in the Great Basin Desert, likely while colonizing new desert habitat that became available as climates warmed following the Last Glacial Maximum.

Gut microbiomes can change quickly and predictably alongside host evolution, implying that host-microbe interactions are important drivers of host adaptation and diversification [50, 51]. As such, we propose that scorpion venom microbiomes may have diverged concurrently with host clade formation during the Pliocene and Pleistocene. Furthermore, the increased diversity in Great Basin Desert samples may reflect a phenomenon where venom microbiomes become more diverse as their hosts colonize and adapt to novel habitats.

As a species expands its range into new environments, individuals encounter and acquire novel environmental microbes through various routes e.g., [52, 53], which can lead to an enrichment of their internal microbial communities. This increased diversity may confer an adaptive advantage, as a more diverse microbiome can offer “colonization resistance” against opportunistic or pathogenic microbes e.g., [54, 55], which could be particularly beneficial in challenging or unfamiliar environments like the newly formed Great Basin Desert. Furthermore, host adaptation to new habitats can involve co-evolutionary processes with their microbiomes, where selective pressures favor symbionts that expand the host’s abiotic niche and enable colonization of otherwise harsh environments, a process known as biotic facilitation (reviewed in [56]). Thus, the proposed recent, potentially postglacial, range expansions of *P. becki* and *A. phaiodactylus*, as well as other co-distributed Great Basin Desert taxa [57–59], provide a temporal framework for such a diversification of their associated venom microbiomes.

In conclusion, we have observed a partial convergence of certain microbial taxa across diverse venomous organisms, independent of habitat and physiology. This pattern suggests that venom secretions may represent a unique and underexplored microbial niche influenced by both environmental factors and strong venom-derived selective pressures. Further comparative studies across scorpion populations and habitats is warranted to shed more light on how geographic distribution and ecological behavior may influence the composition of microbial communities.

## Acknowledgments

We thank our former undergraduate students, Lauren Atkinson, Maxime Parent, and Christopher Shimwell, for initiating the preliminary experiments that resulted in this comprehensive study. Paula Cushing assisted with collecting permits. Funding was provided by the Connecticut State University-American Association of University Professors (CSU-AAUP; BM, CESL, MRG).

## Supporting information captions

**S1 Fig. Principal Component Analysis plots of *P. becki* microbiome samples from exterior surface swabs and venom.** Ellipses represent 95% confidence intervals for the grouped distance metrics (PERMANOVA, 999 permutations, Bray Curtis, p=0.006; Jaccard, p=0.004; Weighted Unifrac, p=0.036; Unweighted Unifrac p = 0.118).

**S2 Fig. Rarefaction curves of venom-associated microbiome samples from *A. phaiodactylus* and *P. becki* regarding ASVs**. Vertical purple dashed line represents the rarefying sequence depth used for downstream analysis.

**S3 Fig. Venn diagram of ASV presence/absence for A*. phaiodactylus* and *P. becki*venom microbiome**. All ASVs at 2,100 rarefaction level.

**S4 Fig. Heatmap scores for the predicted metabolic pathways from PICRUSt2 analysis that function in the venom microbiome of *A. phaiodactylus* vs *P. becki***.

## Tables

**S1 Table. The number of core microbiome ASVs and their fraction of the total number of ASVs identified at different levels of occurrence in *A. phaiodactylus* (n=31) and *P. becki* (n=23) samples**.

**S2 Table. Pairwise Kruskal-Wallis statistical test for alpha diversity metrics between *Anuroctonus phaiodactylus* (n=31) and *Paruroctonus becki* (n=23). Input samples were rarefied at a depth of 2,100 sequences**.

